# Dietary, Cultural and Pathogens-related Selective Pressures Shaped Differential Adaptive Evolution Among Native Mexican Populations

**DOI:** 10.1101/2021.04.14.439124

**Authors:** Claudia Ojeda-Granados, Paolo Abondio, Alice Setti, Stefania Sarno, Guido Alberto Gnecchi-Ruscone, Eduardo González-Orozco, Sara De Fanti, Andres Jiménez-Kaufmann, Héctor Rangel-Villalobos, Andrés Moreno-Estrada, Marco Sazzini

## Abstract

Native American genetic ancestry has been remarkably implicated with increased risk of diverse health issues in several Mexican populations, especially in relation to the dramatic changes in environmental, dietary and cultural settings they have recently undergone. In particular, the effects of these ecological transitions and Westernization of lifestyles have been investigated so far predominantly on Admixed individuals. Nevertheless, indigenous groups, rather than admixed Mexicans, have plausibly retained the highest proportions of genetic components shaped by natural selection in response to the ancient milieu experienced by Mexican ancestors during their pre-Columbian evolutionary history. These formerly adaptive alleles/haplotypes have the potential to represent the genetic determinants of some biological traits peculiar to the Mexican people and a reservoir of loci with potential biomedical relevance. To test such a hypothesis, we used high-resolution genomic data to infer the unique adaptive evolution of 15 Native Mexican groups selected as reasonable descendants of the main pre-Columbian Mexican civilizations. A combination of haplotype-based and gene-network analyses enabled us to detect genomic signatures ascribable to polygenic adaptive traits evolved by the main genetic clusters of indigenous Mexican populations to cope with local environmental and/or cultural conditions. Some of them were also found to play a role in modulating the susceptibility/resistance of these groups to certain pathological conditions, thus providing new evidence for diverse selective pressures having contributed to shape current biological and disease-risk patterns in present-day Native and Mestizo Mexican populations.

## Introduction

Mexican populations represent a paradigmatic example of human groups that have recently experienced dramatic shifts in their environmental, dietary and cultural settings, which are expected to have had relevant social and biological consequences, especially for those communities that have still maintained high proportions of Native American (NA) ancestry. In fact, NA genetic background has been implicated with increased risk of a variety of health outcomes in Mestizo (i.e. Admixed) individuals, in whom the effects of recent ecological transitions and Westernization of lifestyles have been predominantly investigated (Rivera et al. 2004; Aguilar-Salinas et al. 2011; Weissglas-Volkov et al. 2013; Ko et al. 2014).

Conversely, few studies have focused so far on Native Mexican populations (Acuña-Alonzo et al. 2010; Lara-Riegos et al. 2015; Romero-Hidalgo et al. 2017), who are instead the most suitable groups to be considered for inferring the pre-Columbian evolutionary history of the ancestors of modern Mexican people (Moreno-Estrada et al. 2014; Ávila-Arcos et al. 2020). Interestingly, these studies revealed a complex genetic structure of the main indigenous groups that have long-inhabited the present-day Mexico’s territory, which is tightly related to the well-known historical, cultural and demographic processes they have experienced (Moreno-Estrada et al. 2014; Romero-Hidalgo et al. 2017; Ávila-Arcos et al. 2020). In particular, a northwest-southeast gradient of Mexican variation has been described, along with an overall differentiation in the proportions of ancestry components between northern, central and southern populations, which is clearly distinguishable in both Native and geographically-neighboring Mexican Mestizo groups (Rubi-Castellanos et al. 2009; Moreno-Estrada et al. 2014). Major genetic divergence between Aridoamerican (i.e. northern) and Mesoamerican (i.e. central/southern) indigenous populations has been previously suggested to be ascribable also to the contrasting climate and ecological conditions characterizing these Mexican macro-regions (Gorostiza et al. 2012; González-Martin et al. 2015). Nonetheless, the adaptive evolution of Native Mexican ancestors in response to local environmental and/or cultural selective pressures has been scarcely investigated so far (Ávila-Arcos et al. 2020). Unfortunately, this has limited also the identification of the genetic determinants of important biological features of present-day Native and Mestizo Mexican people. In particular, variation underlying adaptive traits that have been shaped by natural selection in relation to the ancestral milieu of these human groups might represent an important reservoir of alleles/haplotypes with potential biomedical relevance (Sazzini et al. 2016; Ávila-Arcos et al. 2020; Sazzini et al. 2020; Landini et al. 2020). In fact, the rapid and substantial modifications occurred in modern Mexican societies might have turned some of these biological adaptations into unfavorable (i.e. dis-adaptive) traits, thus contributing to differential disease susceptibility among populations with varying proportions of NA ancestry. To test such a hypothesis, we used high-resolution genomic data to infer the adaptive evolution of 15 Native Mexican groups selected as reasonable descendants of the main pre-Columbian Mexican civilizations. For this purpose, we imputed genome-wide data from the Native Mexican Diversity Panel (NMDP) (Moreno-Estrada et al. 2014) and we applied a combination of haplotype-based and gene-network analyses to detect genomic signatures ascribable to the occurrence of selective events under a realistic approximation of a polygenic adaptation model (Gnecchi-Ruscone et al. 2018; Sazzini et al. 2020). Accordingly, we identified different adaptive traits peculiar to the main genetic clusters of Native Mexican populations, thus providing new evidence for diverse selective pressures having contributed to shape their current biological and disease-risk patterns.

## Results

After genotypes imputation of the NMDP and multiple pre- and post-imputation quality control (QC) filtering (see Materials and Methods), we obtained a “high-quality imputed dataset” including 4,875,751 single nucleotide variants (SNVs) characterized for 271 individuals from 15 Native Mexican populations (supplementary table S1, Supplementary Material online). These subjects were selected according to their high proportions of NA ancestry based on results from ADMIXTURE analysis (fig. 1, and supplementary fig. S1, Supplementary Material online).

**Fig. 1.**
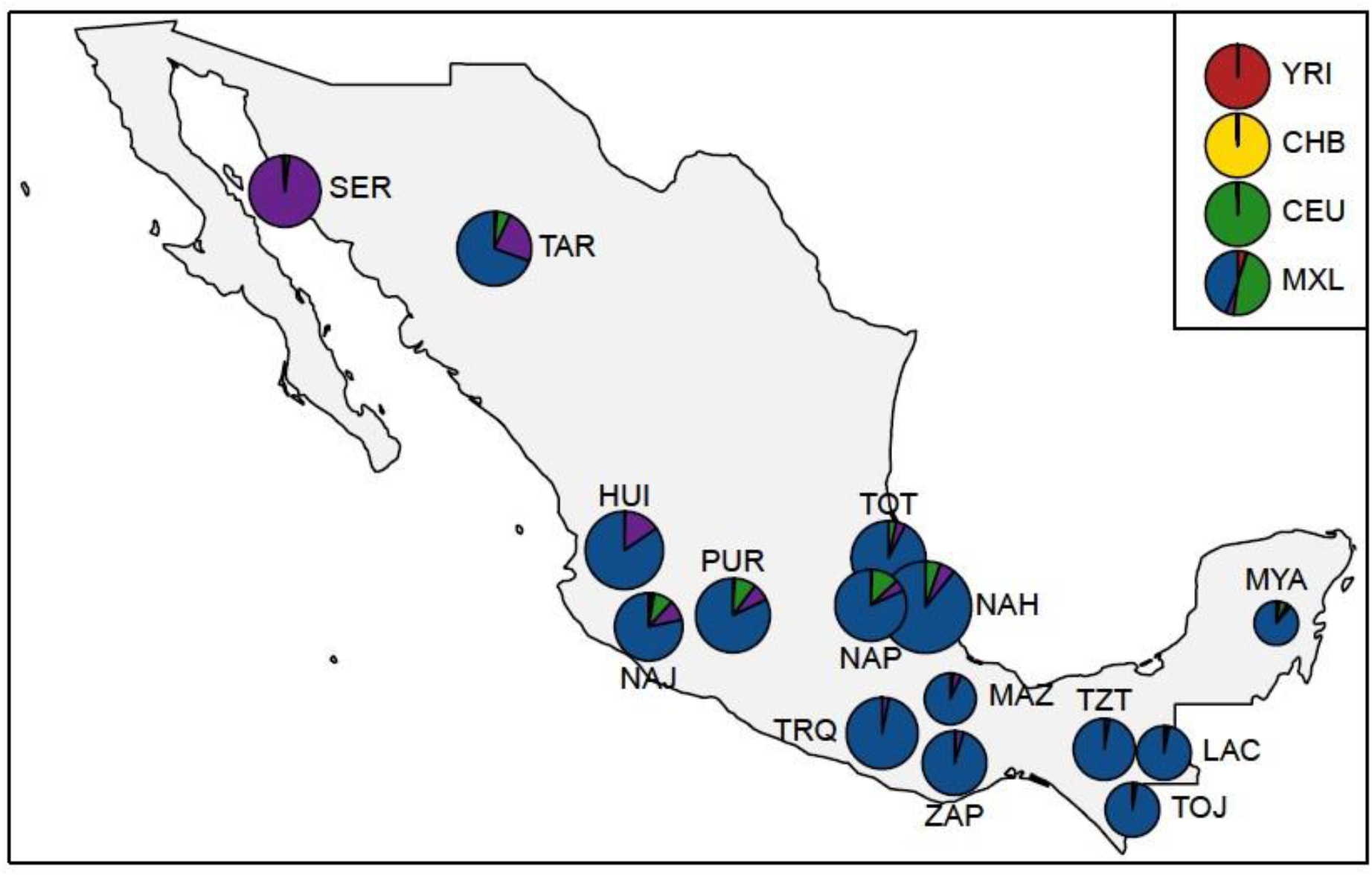
Average proportions of ancestry components inferred for the 15 Mexican indigenous groups included in the NMDP by means of ADMIXTURE unsupervised clustering analysis at K = 5. Evaluation of individual ancestry profiles were used to select subjects representative of the genomic background of their population of origin and showing predominant proportions of Native American ancestry (in dark blue). Individuals of African (YRI), East Asian (CHB), and European (CEU) ancestry, as well as Admixed Mexican-American samples (MXL), were also included to quantify non-Native American genetic fractions observable in the indigenous genomes. SER showed a peculiar ancestry component (in purple) whose high proportion was plausibly ascribable to strong genetic drift experienced due to prolonged isolation, as previously attested (Moreno-Estrada et al. 2014). Pie charts diameters are proportional to the sample sizes of the considered groups (details for each population, full set of K tested and cross-validation errors are reported in supplementary table S1 and supplementary fig. S1, Supplementary Material online). The geographical map has been plotted using the R software (v.4.0.2).

To test the concordance of the imputed data with patterns of genomic variation previously described for the examined populations, we compared results obtained by means of Principal Component Analyses (PCA) performed before (Moreno-Estrada et al. 2014) and after the imputation procedure (supplementary fig. S2, Supplementary Material online). In both cases, the well-known northwest to southeast cline of Mexican genetic variation was observed (supplementary fig. S2, Supplementary Material online).

### Depicting Patterns of Haplotype Sharing Among Native Mexican Populations

We then explored the fine-scale genetic structure of these populations by applying the haplotype-sharing clustering approach implemented in the CHROMOPAINTER/fineSTRUCTURE pipeline. This first enabled us to remove six outlier individuals that did not show close genetic affinity with those belonging to their population of origin and may be thus representative of misleading mismatches between their self-reported and genomic ancestry. By considering only branches of the obtained fineSTRUCTURE dendrogram showing posterior probability above 90%, we then identified three main clusters of Native Mexican populations: the Northern Mexican cluster (NMC), the Central Mexican cluster (CMC), and the Southern Mexican cluster (SMC) (fig. 2). This finding was already attested by previous studies (Gorostiza et al. 2012; Moreno-Estrada et al. 2014), but our fineSTRUCTURE analysis pinpointed a different distribution of some groups within the identified clusters. In detail, the NMC included Seri (SER) and Rarámuri (also known as Tarahumara, TAR) populations from northern Mexico. The CMC was made up of people from Central West and Central East Mexico and belonging to the Huichol (HUI), Nahua (NAH), Nahua Jalisco (NAJ), Nahua Puebla (NAP) and Purepecha (PUR) populations. The SMC instead grouped populations from South (i.e. Triqui, TRQ; Mazatec, MAZ; Zapotec, ZAP) and South East (i.e. Tojolabal, TOJ; Tzotzil, TZT; Lacandon, LAC; Maya, MYA) Mexico, also integrating the Totonaca (TOT) from Central East Mexico (fig. 2). In particular, an appreciable genetic differentiation was observed between NMC populations and CMC or SMC ones (F_st_ = 0.023, p < 10^−6^; F_st_ = 0.029, p < 10^−6^, respectively), while CMC and SMC turned out to be considerably less differentiated (F_st_ = 0.007, p < 10^−6^). However, when considering NMC populations, a significant differentiation between SER and TAR (F_st_ = 0.074, p < 10^−6^) was also evident.

**Fig. 2.**
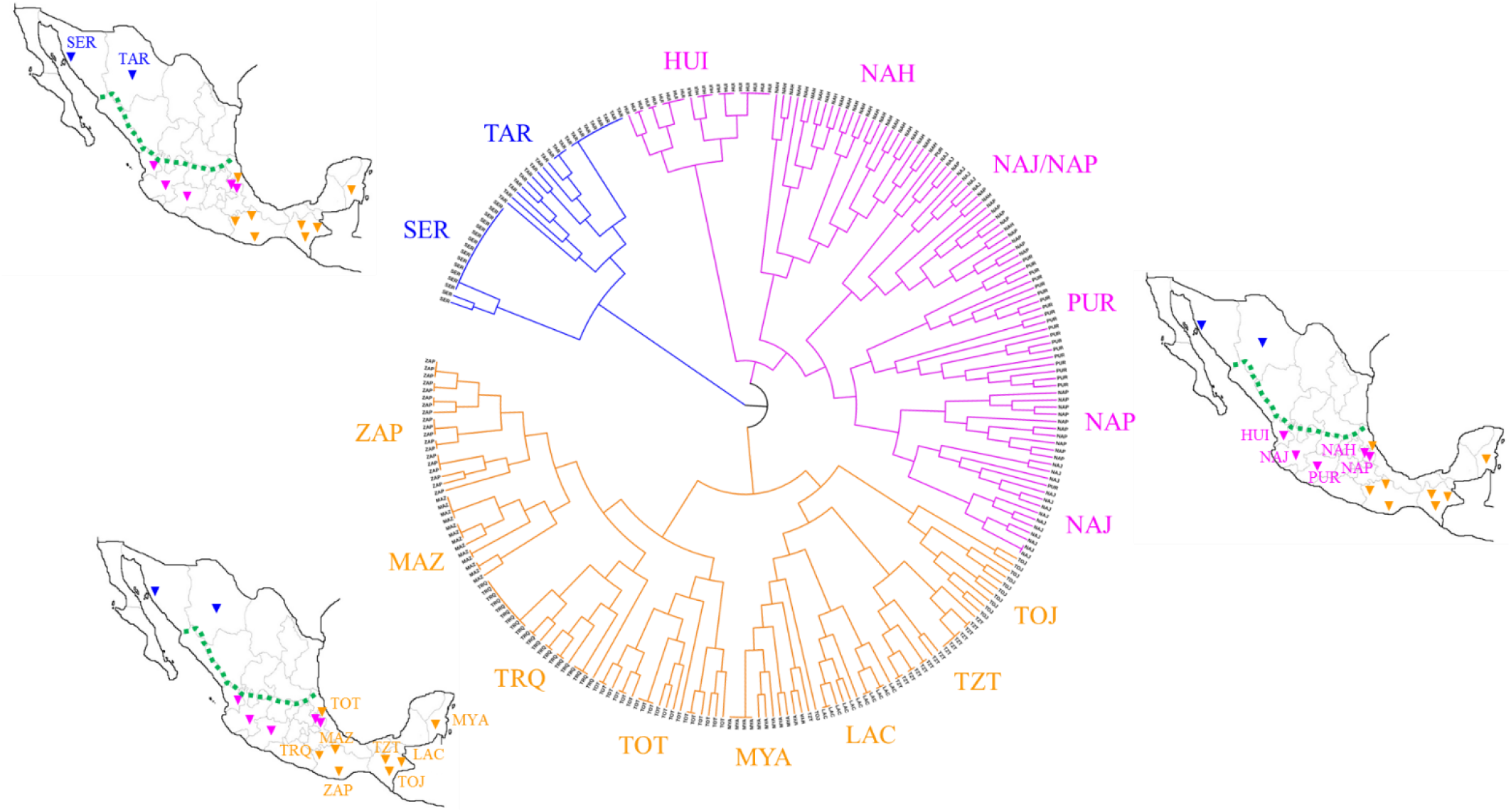
fineSTRUCTURE hierarchical clustering dendrogram based on haplotype sharing patterns observed among the considered Native Mexican groups and their geographical locations. Population clusters were defined by considering branches of the dendrogram that split with a posterior probability above 90%. Accordingly, the Northern Mexican Cluster (in blue) included SER and TAR populations, the Central Mexican Cluster (in fuchsia) was made up of groups from Central West and Central East Mexico (i.e. HUI, NAH, NAJ, NAP, and PUR), while the Southern Mexican Cluster (in orange) grouped populations from Southern Mexico (i.e. TRQ, MAZ, ZAP), South East Mexico (i.e. TOJ, TZT, LAC, MYA), and Central East Mexico (i.e. TOT). The dotted green line indicates the demarcation between Aridoamerican (northward) and Mesoamerican (southward) regions, which represent the main geographical and cultural areas of ancient Mexico and whose population clusters showed the greatest genetic differentiation according to F_st_ analyses. The geographical maps have been plotted using the R software (v.4.0.2).

### Differential Genomic Signatures of Positive Selection Across Native Mexican Clusters

We then computed the Number of Segregating sites by Length (nSL) statistics and we used the obtained nSL genome-wide distributions to perform gene-network analyses (see Materials and Methods) in order to infer the adaptive evolution of each cluster by assuming that populations belonging to the same genetic group have presumably experienced similar environmental and/or cultural selective pressures. The sole exception was represented by NMC populations, who were analyzed separately due to their significant genetic differentiation and because they have long maintained appreciably different lifestyles despite having encountered similar environmental conditions in Aridoamerica (MacWilliams et al. 2008; Rentería Valencia 2009).

#### Seri Population

Results obtained by selection scans seems to be consistent with the above-mentioned hypothesis. In detail, three gene networks made up of 18 genes were found to be significantly enriched for selective events in the SER population (fig. 3a, supplementary table S2, Supplementary Material online). Seven of these genes are known to be implicated in the *Starch and sucrose metabolic pathway* and, according to the Kyoto Encyclopedia of Genes and Genomes (KEGG) database, they also play a role in the *Galactose metabolism* and *Fructose and mannose metabolism* by contributing to the degradation/utilization of other simple sugars. Moreover, they also participate in the regulatory steps of the *Glycolysis/Gluconeogenesis* and *Insulin signaling pathway.* In particular, the *SI* and *TREH* genes are highly expressed in the brush border of the small intestine, where the final stage of carbohydrate digestion occurs, including that of sucrose, trehalose and starch intermediates (UniProt Consortium 2019). Conversely, the *HK2, HK1, HKDC* and *GCK* genes encode proteins of the hexokinase family that catalyze the phosphorylation of hexoses to produce D-fructose-6-phosphate or D-glucose-6-phosphate, the first and ratelimiting step in most glucose metabolism pathways (UniProt Consortium 2019). Interestingly, as reported in the KEGG database, these latter loci are also involved in the pathogenesis of type II diabetes.

**Fig. 3.**
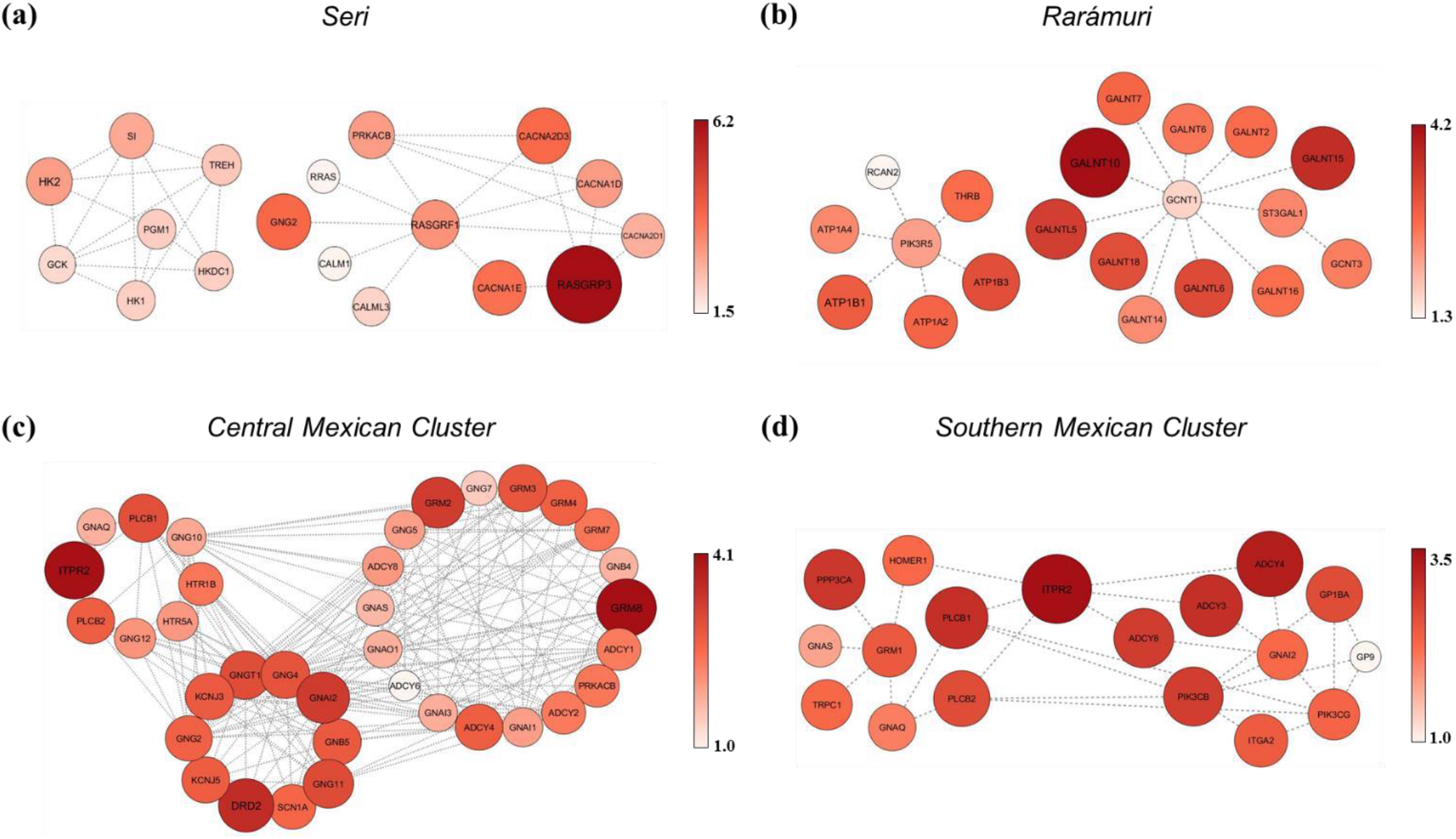
Representation of gene networks mediating polygenic adaptive events inferred for Seri, Rarámuri and genetic clusters of Native Mexican populations identified according to fineSTRUCTURE analysis. (a) Gene networks belonging to the *Starch and sucrose metabolism, Ras signaling* and *MAPK signaling* pathways showing significant footprints of positive selection in the SER population. (b) Gene networks belonging to the *Thyroid hormone signaling* and *Mucin type O-Glycan biosynthesis* pathways having experienced multiple selective events in the TAR population. (c) Highly interrelated gene networks belonging to the *Glutamatergic*, *Serotonergic* and *Dopaminergic synapse* pathways showing widespread selection signatures in Central Mexican Cluster populations. (d) Gene networks belonging to the *Glutamatergic synapse* and *Platelet activation* pathways having mediated adaptive evolution of Southern Mexican Cluster populations. Color intensity of circles is displayed according to the reported color-scale and, along with circles size, is proportional to the *per gene* nSL representative scores used to perform gene-network analyses.

The other two significant gene networks observed in the SER population included 11 genes primarily involved in the *Ras signaling* and *MAPK signaling* pathways, which turned out to be functionally related to the *RASGRF1* gene (fig. 3a, supplementary table S2, Supplementary Material online). Signaling through the Ras/Raf/MAPK cascade responds primarily to growth factors and mitogens to ultimately regulate cell proliferation, differentiation, and apoptosis (Morrison 2012). The loci that showed the most relevant selection signatures in these gene networks were *RASGRP3* and *RASGRF1.* They encode for guanine nucleotide exchange factors that activate Ras proteins by switching its Ras-bound GDP to GTP, *GNG2* encoding the gamma subunit of heterodimeric G proteins and acting as an upstream regulator of *RASGRP3* and *RASGRF1*. Moreover, *CACNA2D3, CACNA1E* and *CACNA1D* encode for subunits of calcium voltage-gated channels involved in calcium-dependent processes, such as muscle contraction, hormone or neurotransmitter release, gene expression, cell motility, cell division and apoptosis (UniProt Consortium 2019) (fig. 3a, supplementary table S2, Supplementary Material online).

#### Rarámuri Population

Two gene networks belonging to the *Thyroid hormone signaling* and *Mucin type O-Glycan biosynthesis* pathways and including 20 genes were found to have plausibly experienced multiple selective events in the TAR population (fig. 3b, supplementary table S3, Supplementary Material online).

The thyroid hormones (TH) T3 (triiodothyronine) and T4 (thyroxine) play an important role in the organism’s growth and development, as well as in the regulation of glucose and lipid metabolism, thermogenesis and energy homeostasis, with T3 being considered the main active form and T4 instead its precursor. Interestingly, the identified putative adaptive genes are primarily involved in the alternative non-canonical and more rapid pathway of TH and encode for Na^+^/K^+^-ATPases (e.g. *ATP1B3, ATP1B1, ATP1A2* and *ATP1A4),* TRβ isoforms *(THRB),* a subunit of PI3K proteins *(PIK3R5),* and a calcineurin inhibitor *(RCAN2)* (fig. 3b) (Moeller and Broecker-Preuss 2011). Through the non-canonical actions of TH, regulation of gene expression is achieved by an initial non-transcriptionally mediated change in cell physiology. For example, TH can induce modifications in the phosphorylation and activation patterns of the PI3K pathway, as well as in the modulation of availability of second messengers and of plasma membrane ion pump activity (e.g. Na^+^/K^+^-ATPase, Na^+^/H^+^ exchanger and Ca^++^-ATPase) (Cordeiro et al. 2013; Senese et al. 2014). These alternative mechanisms play a relevant role in energy metabolism by regulating glucose and triglyceride concentrations, body temperature, locomotor activity, and heart rate (Hönes et al. 2017). Indeed, TRβ isoforms *(THRB)* are expressed mainly in cardiac and skeletal muscle, brain and adipose tissue.

Candidate genes belonging to the *Mucin type O-Glycan biosynthesis* pathway instead participate to *O*glycosilation of mucins, a posttranslational modification mediated by glycosyltransferase enzymes and consisting in the addition of monosaccharides (*O*-glycans) to the hydroxyl group of serine or threonine residues. Mucins represent a family of large and heavily N-acetylgalactosamine (*O*-GalNAc) glycosylated proteins highly expressed by epithelial tissues that produce gel-like secretions to lubricate and protect airways, urogenital and gastrointestinal tracts against physical, chemical or biological insults. Mucin-type *O*-glycans can also serve as surface receptors for adhesion molecules, mediate the interaction with pathogens including bacterial, fungal and viral infections, and locally modulate the inflammatory response (Bergstrom and Xia 2013; Duarte et al. 2016). Among *O*-glycans, *O*-GalNAc is especially relevant for mucin formation and genes showing selection signatures in the TAR population encode for several GalNac transferases (*GALNT* genes) that fine-tune mucin biosynthesis (fig. 3b).

#### Populations Belonging to the Central Mexican Cluster

As regards CMC populations, gene networks belonging to the *Glutamatergic*, *Serotonergic* and *Dopaminergic synapse pathways* showed widespread signatures of positive selection distributed across 36 highly interrelated and overlapping genes mediating dependence on addictive substances such as alcohol (fig. 3c, supplementary table S4, Supplementary Material online). Glutamate, serotonin (5-Hydroxytryptamine, 5-HT) and dopamine are indeed important neurotransmitters in the central nervous system, with glutamate representing the major excitatory one and being involved in synaptic plasticity, memory retention and learning, while serotonin regulates behavioral and physiological functions, such as sleep and wake states, emotions, and hemostasis, among others. Dopamine, instead, controls locomotor and visceral functions, motivational and rewarding behaviors, and has been implicated in addiction to activities and substances providing excitement and pleasure (Kanehisa and Goto 2000; Niyonambaza et al. 2019). Interestingly, genes showing some of the most relevant selection signatures in CMC populations encode proteins that modulate inhibitory mechanisms of these systems by suppressing the excess release of neurotransmitters. In detail, Group II *(GRM2, GRM3)* and III *(GRM8, GRM4* and *GRM7*) metabotropic glutamate receptors (mGluRs), the serotonin 5-HT1B *(HTR1B)* receptor, and the dopamine receptor D2 (*DRD2*) are Gi/o type G-protein coupled autoreceptors. They are known to suppress excess glutamate, 5-HT, and dopamine release via inactivation of the adenylate cyclase/cAMP/protein kinase pathway and the activation of G-protein-coupled inwardly rectifier K^+^ channels-GIRKs (*KCNJ3* and *KCNJ5*), leading to membrane hyperpolarization and suppression of neuronal excitability. In addition, the splice variant of the G-α_i2_ protein *(GNAI2)* regulates the cell surface density of DRD2 by sequestrating them as an intracellular reservoir (López-Aranda et al. 2007). Among the other genes showing smoother selection signatures in these gene sub-networks some encoding the alpha, beta and gamma subunits of G-proteins (*GNAIs*, *GNAs*, *GNBs* and *GNGs),* the members of the adenylate cyclase family *(ADCYs)* and a member of the protein kinase cAMP family (*PRKACB*) were observed. *ITPR2*, *PLCB1* and *PLCB2* genes are instead specifically involved in mobilization of intracellular calcium (Ca^2+^) stores. However, the mechanism of action of the named neurotransmitters, especially their excitatory neurotransmission, also requires intracellular Ca^2+^ mobilization via their coupling to G-proteins that activates phospholipases C *(PLCBs)* to form inositol 1,4,5-trisphosphate (IP3). IP3 then binds to the IP3Rcalcium release channels receptors (*ITPR2*) in the endoplasmic reticulum to increase eventually cytosolic Ca^2+^ levels. When cross-checking the identified putative adaptive loci with information from the KEGG database, genes belonging to these pathways, which are expressed ubiquitously in diverse tissues, were found to contribute the *Alcoholism*, *Morphine addiction*, *Retrograde endocannabinoid signaling* and *Cocaine addiction* pathways. Moreover, they were found to play a role also in immunological and inflammatory responses to viruses and trypanosomes by participating in the *Human cytomegalovirus infection*, *Chagas disease* and *Chemokin signaling* pathways.

#### Populations Belonging to the Southern Mexican Cluster

SMC populations showed signatures of positive selection very similar to those observed for CMC groups, particularly at genes implicated in immune and inflammatory responses to viruses and trypanosomes (e.g. *ITPR2, ADCY4, ADCY8, PLCB1, PLCB2, GNAI2, GNAQ* and *GNAS*) (fig. 3d, supplementary table S5, Supplementary Material online). In fact, 18 candidate loci were found to participate in the *Glutamatergic synapse* and *Platelet activation* pathways and were characterized by tight functional connections with the *ITPR2* gene, which showed the highest *per gene* nSL value among those computed. In particular, when we framed these putative adaptive loci in the functional cascades annotated in the KEGG database they turned out to be involved also in the *Chemokin signaling, Inflammatory mediator regulation of TRP channels, Human cytomegalovirus infection* and *Chagas disease (American trypanosomiasis)* pathways, thus confirming their primary role in immunity and inflammation. In fact, although platelets have been long known to act as regulators of hemostasis and thrombosis, their additional role in innate immune reactions against bacteria, viruses and even tumors has been recently demonstrated. Indeed, they have shown to express nine Toll-like receptors differently by gender during pathogen detection, as well as to have the ability to sample the blood environment to present viruses or bacteria to other immune cells (Koupenova et al. 2015; Holinstat 2017). Interestingly, functioning of the *Platelet activation pathway*, including granule secretion and spreading, is known to depend on the increase of intracellular Ca^2+^ concentration, and indeed *ITPR2*, *PLCB1* and *PLCB2* loci, which are found to be putative adaptive genes also in the CMC, are particularly implicated in the mobilization of intracellular Ca^2+^ stores.

## Discussion

To address the lack of studies specifically aimed at inferring the adaptive evolution of Native Mexican human groups, we focused on populations selected as reasonable descendants of the main pre-Columbian civilizations that inhabited the geographical regions nowadays attributed to the Mexican territory. For this purpose, we used the NMDP to collect data for groups showing high proportions of NA ancestry and representative of the main gene pools distinguishable within the overall Mexican population (fig. 1) (Moreno-Estrada et al. 2014; Ávila-Arcos et al. 2020). However, previous studies conducted on these indigenous populations relied on SNP-chip data and/or exome variation, which prevented the exploration of the full spectrum of Mexican alleles in terms of both frequency and nature, thus limiting the potential in investigating fine-scale patterns of population structure and polygenic adaptive processes. In the attempt to overcome this issue, we imputed genome-wide genotypes from the NMDP to considerably increase the resolution of the available genomic dataset.

### Fine-scale Genetic Structure of Native Mexican Populations

FineSTRUCTURE haplotype-sharing clustering analysis first enabled us to identify three broad groups of genetically homogenous Native Mexican populations that are expected to have shared a remarkable fraction of their evolutionary histories (fig. 2). The observed patterns of fine-scale population structure were found to be consistent with the putative ancestral geographic location of the bulk of populations that composed these genetic clusters (fig. 1, supplementary fig. S2, Supplementary Material online). The TOT group from Central East Mexico represented the sole exception, being settled near the Gulf Coast, but clustering within the SMC. This peculiar outcome was likely ascribable to the presence of a geographic barrier (i.e. the Sierra Madre Oriental) that segregated TOT from the other CMC groups. Moreover, TOT are known to have historically maintained hostile relationships with neighboring populations such as the Aztecs, who represented the ancestors of present-day Nahua people, instead establishing intense social and commercial exchanges with southern Mexican populations (Báez-Jorge 2009). Such a genetic affinity between TOT and SMC groups is consistent also with osteological, archeological, and ethnohistorical evidence (Ragsdale and Edgar 2015) and is in agreement with previous findings pointing to gene flow between TOT and MYA people from South East Mexico (Moreno-Estrada et al. 2014).

Overall, a major genetic differentiation was observed between NMC and CMC or SMC (F_st_ ≥ 0.023, p < 10^−6^) than between CMC and SMC (F_st_ = 0.007, p < 10^−6^). This suggests a main divergence between populations that have inhabited Aridoamerica and Mesoamerica, the two major geographical and cultural areas of ancient Mexico, which have been long characterized by contrasting climates and biodiversity landscapes. Such a fragmented demographic history of Mexican groups was proposed also by previous studies, which pointed to a genetic structure of Native Mexicans that is more related to their territories’ orography rather than to cultural/linguistic differences (Gorostiza et al. 2012; González-Martín and Gorostiza 2013; Moreno-Estrada et al. 2014; Rangel-Villalobos et al. 2016). In particular, our findings are consistent with results from the analysis of exome sequence data of Mexican indigenous populations, including TAR, HUI, TRQ and MYA, which inferred that northern and southern ancestral Native Mexican groups split approximately 7.2 thousand years ago (KYA) and further diverged locally around 6.5 and 5.7 KYA, respectively (Ávila-Arcos et al. 2020). Moreover, more recent differentiation between HUI/PUR (3.4 KYA), NAH/PUR (2.2 KYA), and HUI/NAH (357 YA) seems to support our assumption that these CMC populations largely shared a common evolutionary history since they diverged from NMC and SMC groups (González-Martín and Gorostiza 2013). This pattern is also consistent with the frequency distribution of the four “Pan-American” mtDNA haplogroups (i.e. A2, B2, C1, D1) in Aridoamerican (e.g. Pima) and Mesoamerican indigenous groups (among which HUI, NAH, Otomí, MYA). In fact, geographic discontinuities in their frequencies revealed two major genetic boundaries deeply delimiting Aridoamericans from all Mesoamerican groups, as well as a southern boundary differentiating MYA from central Mesoamerican populations (Gorostiza et al. 2012).

Interestingly, when considering NMC populations, a noteworthy differentiation was observed also between SER and TAR (F_st_ = 0.074, p < 10^−6^) in accordance with the lack of evidence for their historical relationships. SER has indeed shown remarkable linguistic and genetic differentiation with respect to other Native Mexican populations, including TAR, and an independent ancestry component (fig.1, supplementary fig. S1, Supplementary Material online) suggesting a high degree of genetic drift due to prolonged isolation (Moreno-Estrada et al. 2014). In fact, regions inhabited by SER and TAR are separated by a geographic barrier represented by the Sierra Madre Occidental and in addition, among Aridoamericans, SER maintained a semi-nomadic hunter-gatherer lifestyle until the mid-20^th^ century, while there is evidence for TAR ancestors having become mobile farmers since around 1,600 BC (MacWilliams et al. 2008; Rentería Valencia 2009). Therefore, we can speculate that these populations clustered together into the NMC because they were substantially differentiated from both CMC and SMC ones, rather than because they present a considerable genetic affinity.

Based on this body of evidence, we assumed that SER, TAR, CMC and SMC populations might have adaptively evolved independently in response to differential environmental- and/or cultural-related selective pressures. Accordingly, we searched for genomic signatures left by the action of natural selection on these main Native Mexican groups by testing the occurrence of selective events under a realistic approximation of a polygenic adaptation model.

### Seri Metabolic Adaptation to a Diet Rich in Simple Carbohydrates

Metabolism of simple carbohydrates (i.e. mono and disaccharides) and regulation of glucose homeostasis via the *Glycolysis/Gluconeogenesis* and *Insulin signaling* pathways emerged as the biological functions that most plausibly evolved adaptively in the SER population (fig. 3a, supplementary table S2, Supplementary Material online). In fact, selective events at genes playing a role in these metabolic processes seems to be highly consistent with the peculiar diet and lifestyle of this Native Mexican group, who has long-relied mostly on the consumption of fruits of succulent plants that are rich in simple sugars.

According to archeological evidence, this semi-nomadic population originally known as Comcáac has indeed inhabited the dry desert, coastal margins and neighboring islands (e.g. the Tiburón and San Esteban islands in the Gulf of California) of the current state of Sonora in Northwestern Mexico since at least 2 KYA. Moreover, they are known to have maintained their hunting-gathering and fishing based lifestyle until about the 1950s (Rentería Valencia 2009). In particular, they used to organize themselves into mobile groups or “bands”, moving around in camps on the shore and islands during the dry-cold season (September-May), while occupying the desert areas during the wet-hot period (June-August). During the cold season, their diet was based on the tender stem of two species of cholla cacti (i.e. *Cylindropuntia bigelovii* and *C. fulgida),* which provided up to 80% of the total carbohydrates consumed, as well as on hunting products and sea resources (Álvarez Palma and Casiano 2003; Hernández Santana 2017). Instead, during the hot season they consumed mainly fruits and seeds of various species of columnar cacti, among which the saguaro *(Carnegiea gigantea),* sweet pitaya *(Stenocereus thurberi)*, cardon (*Pachycereus pringlei*), sour pitahaya (*Stenocereus gummosus*) and cina cactus (*Stenocereus alamosensis*), from which they also prepared sweet fermented drinks. The sweet pods from mesquite tree and the prickly pears from the cholla and nopal *(Opuntia ficus-indica)* were also highly appreciated during this season (Álvarez Palma and Casiano 2003; Hernández Santana 2017). Therefore, selective events observed at multiple loci contributing mainly to sucrose, fructose and mannose metabolism might have optimized the utilization of the high amounts of ingested sugars as energy sources or reserves, also reducing the potential metabolic risk associated with glucose overload (Jang et al. 2018; Rabbani and Thornalley 2019). In line with such a hypothesis, among the top candidate adaptive genes identified (supplementary table S2, Supplementary Material online), *HK2*, *HK1* and *HKDC* encode for hexokinases that have a high affinity for glucose and in particular *HK2,* which is highly expressed in the skeletal muscle and adipose tissue, controls the rate-limiting step of glycolysis (Tan and Miyamoto 2015). In contrast, *GCK* is known to regulate the rate-limiting step of glycolysis and glycogenesis in pancreatic beta cells and hepatocytes and its product shows weak affinity for glucose, but becomes effective when glucose is abundant by modulating insulin secretion and storage of glucose excess (UniProt Consortium 2019). Furthermore, the SI enzyme was proven to be critically involved in the final digestion step of sucrose and starch intermediates such as isomaltose, while TREH hydrolyzes the disaccharide trehalosa, which is naturally found in plants, algae, insects and shrimp (Elbein et al. 2003; UniProt Consortium 2019). Other putative targets of selection in SER people, which support the above-mentioned hypothesis, included genes involved in signal transduction cascades (i.e. *Ras* and *MAPK signaling* pathways). In particular, they play a role in the signaling cascade leading to insulin secretion, as is the case of *CACNA1E* and *CACNA1D* genes that are also expressed by pancreatic beta cells in the islets of Langerhans (Vajna et al. 1998). Interestingly, these latter loci, as well as *HK2, HK1, HKDC* and *GCK,* have been also proved to be involved in the pathogenesis of type 2 diabetes (Muller et al. 2007; Reinbothe et al. 2013).

### Semi-nomadic Lifestyle and Exposure to Maize Mycotoxins as the Main Selective Pressures Having Acted on the Rarámuri Population

Non-canonical actions of TH that are particularly linked to the regulation of energy metabolism (Hönes et al. 2017), as well as variation at *GALNT* genes controlling *O*-glycosylation processes during the biosynthesis of mucins (Bergstrom and Xia 2013), were inferred to have been shaped by positive selection in the TAR population (fig. 3b, supplementary table S3, Supplementary Material online). These adaptations might have evolved in response to selective pressures due to the peculiar mobile agricultural lifestyle adopted by the ancestors of this Native Mexican group since around 1,600 BC, when they first settled the mountains and canyons of the southern Chihuahua State (Northwestern Mexico) (MacWilliams et al. 2008). Throughout this geographic area of contrasting relief, temperatures (which drop below −20 °C in winter, while rising up to 40 °C in summer), rainfall and vegetation, TAR are indeed known for their traditional residential mobility and reliance on food stored year-round. In fact, over the year they move from their primary residences on the valleys to their growing-season residences (May - June), which are associated with scattered fields for maize cultivation and maize depots, and to their winter locations (December - February) in rock-shelters located in the walls of small canyons. These seasonal shifts of residence require them to routinely walk extended distances with heavy loads from the maize depots to maintain or replenish their supplies (Hard and Merrill 1992). Furthermore, according to archaeological evidence, by 500-1,000 AD, TAR had already devised various unique methods of maize storage, such as underground, cave niches, rectangular and cylindrical structures of stone and cement, or vases of braided grass and clay, thus confirming their strong need to store food for long time as the main subsistence strategy (Hernández Xolocotzi 1985).

It is therefore plausible that the detected selective events might have enhanced two major biological functions that appear to be essential for the TAR population. On the one hand, improvement of the regulation of energy metabolism and thermoregulation might have evolved in response to long-distance walks with heavy loads in such up-and-down relief and extreme climatic conditions. On the other hand, gastrointestinal protection through effective mucins formation could represent a possible adaptation to the ingestion of mycotoxins present in the stored food they relied on. In fact, through the rapid non-canonical signaling mediated by the *THRB*, *ATPases*, and *PIK3R5* putative adaptive genes, TH have been shown to increase energy metabolism, body temperature, oxygen consumption, and locomotor activity, as well as to reduce serum triglycerides concentrations by using them as a substrate for energy production (Hönes et al. 2017; Iwen, Oelkrug, and Brabant 2018). In particular, TRβ^GS^ mice lacking canonical TH receptors signaling, but preserving their non-canonical TH actions, have shown higher VO_2_ (i.e. volume of oxygen uptake) and distance traveled compared to control mice (Hönes et al. 2017). Such a hypothesis of enhanced energy metabolism of TAR is in line with that proposed by two previous studies reporting an enrichment of genes involved in musculoskeletal functions harboring novel missense/promoter region variants (Romero-Hidalgo et al. 2017) and a candidate adaptive gene *(BCL2L13)* highly expressed in skeletal muscle (Ávila-Arcos et al. 2020). Both findings suggested adaptive musculoskeletal traits underlying the well-known physical endurance of people of TAR ancestry.

Finally, studies evaluating the effects of exposure to mycotoxins, such as those frequently found in maize (e.g. deoxynivalenol, zearalenone, fumonisin B1, and aflatoxin B1) in relation to its storage conditions have demonstrated a marked disruption of intestinal epithelial cells and mucus barrier integrity and function (Robert et al. 2017). In detail, the assessment of the effects exerted by deoxynivalenol and fumonisin B1 on mucus has revealed that they modulate both mucin production in the duodenum and its monosaccharide composition (namely of GalNAc, galactose and N-acetyl-neuraminic acid) by regulating the expression of genes involved in mucin biosynthesis (Wan et al. 2014; Pinton et al. 2015). Accordingly, since TAR derive approximately 75% of their diet from maize and have relied mostly on stored maize (Hard and Merrill 1992), it is likely that they have been long exposed to mycotoxin ingestion, which might have represented a selective pressure that led to enhanced mucin biosynthesis aimed at ensuring gastrointestinal homeostasis.

### Adaptation of Central Native Mexican Populations to Addictive Substances

Populations belonging to the CMC were found to have evolved adaptations mostly mediated by functional pathways involved in the dependence on addictive substances, among which alcohol, morphine, and cocaine (fig. 3c, supplementary table S4, Supplementary Material online). Indeed, the most plausible candidate adaptive genes identified in this population group encode autoreceptors for mGluRs (i.e. *GRM2, GRM3, GRM8, GRM4* and *GRM7),* serotonin *(HTR1B)* and dopamine (*DRD2*), which spearhead the inhibitory mechanisms of the related neurotransmission systems and that have been associated with addictive phenotypes (Abrahao, Salinas, and Lovinger 2017; Roberto and Varodayan 2017). These findings seem to be highly consistent with the traditional consumption of alcohol and psychoactive plants by the ancestors of these Native Mexican populations (Roman et al. 2013). Accordingly, we propose that adaptive changes at the *Glutamatergic, Serotonergic* and *Dopaminergic synapse* pathways might have enhanced biological tolerance towards such stimulant compounds and their detrimental effects, which in turn could have led to the development of a phenotype of dependence. In fact, ethno historical evidence suggests that routine consumption of stimulant substances by the ancient Wixárika (HUI) and Aztec (Nahua) populations followed the processes of plants domestications in Mesoamerica approximately 5 KYA. During this period, they also incorporated into their culture the use of fermented alcoholic beverages made from maize *(tejuino)* and the agave plant (“*pulque*”), respectively, which were endemic to this region. In particular, alcoholic beverages were regularly consumed for religious, nutritional, and medicinal purposes, to the extent that excessive drinking leading to drunkenness became strictly prohibited and penalized by death (Roman et al. 2013). Interestingly, most of the identified putative adaptive genes are linked to the ethanol-induced changes in interneuron function and synaptic transmission associated with chronic rather than acute alcohol exposure and play a role in the development of ethanol seeking behavior and alcohol use disorders. For example, neuroadaptations of glutamatergic transmission due to chronic ethanol exposure (CEE) and withdrawal include generally enhanced function of ionotropic (iGluRs) and mGluRs glutamate receptors, resulting in increased release but decreased reuptake of glutamate in several brain regions and an extracellular hyperglutamatergic state (Roberto and Varodayan 2017). In particular, downregulation of mGluR2 encoded by *GRM2* in nucleus accumbens (NAc) neurons contributed to the hyperglutamatergic state and has been implicated in alcohol dependence (Meinhardt et al. 2013). CEE also results in diminished serotonergic and dopaminergic transmission through variable effects on 5-HT receptors, hypoactivity of dopamine neurons, and decreased NAc/striatum dopamine release and DRD2 availability (Abrahao, Salinas, and Lovinger 2017). In addition, *in vivo* studies have revealed that DRD2/GIRK-mediated autoinhibition is uniquely regulated by a Ca^2+^-dependent desensitization mechanism involving IP3-induced Ca^2+^ release and stimulated by mGluRs activation (Perra et al. 2011). In fact, direct molecular targets of ethanol include specific ion channels, such as the GIRKs, encoded by the identified putative adaptive *KCNJ3* and *KCNJ5* genes, as well as non-ion-channel targets, such as intracellular signaling molecules encoded by *ADCYs* and *PRKACB* loci (fig. 3c, supplementary table S4, Supplementary Material online) (Abrahao, Salinas, and Lovinger 2017). Moreover, the *HSD17B11* (a shortchain alcohol dehydrogenase that metabolizes secondary alcohols and ketones) and *KCNC2* genes have been recently identified as targets of natural selection in the HUI population by a study implementing different methodological approaches with respect to the present one (Ávila-Arcos et al. 2020). This further supports the hypothesis of adaptive traits evolved in response to traditional consumption of stimulant substances, particularly alcohol. This seems to be in line also with findings showing that Nahua and HUI populations exhibit one of the highest frequencies documented to date (67% and 65%, respectively) of the *DRD2/ANKK1* TaqIA A1 allele (rs1800497), which is associated with diminished *DRD2* expression and alcohol dependence (Panduro et al. 2017). Interestingly, this variant was found to be not associated with excessive alcohol consumption or metabolic disturbances in these Native Mexican groups. Instead, in the Admixed Mexican population from the same geographic region, the A1 allele was associated with heavy alcohol drinking, overconsumption of palatable foods known to have addictive properties (e.g. meat, fried dishes and sugars), as well as increased risk of impaired glucose, triglycerides, and VLDL levels. Moreover, the C allele of the *DRD2* C957T SNP (rs6277) was associated with increased consumption of sugar and serum triglycerides (Ramos-Lopez et al. 2018; RiveraIñiguez et al. 2019). Although the described genetic signatures overall suggest an ethanol-driven selective pressure, some of the identified putative adaptive genes (i.e. *ITPR2, PLCB1* and *PLCB2),* which were found to have evolved adaptively in SMC populations as well, are also known to regulate intracellular Ca^2+^ mobilization and to be implicated in immunological and inflammatory responses to cytomegalovirus and trypanosomes. In fact, parasites such as the American *Trypanosoma cruzi* and *Leishmania Mexicana* are quite common in the central-southern regions of Mexico and therefore we cannot rule out the hypothesis that also pathogen-related selective pressures have contributed to trigger the inferred adaptive events.

### Adaptations of Southern Native Mexican Populations to Endemic Pathogens

SMC populations were found to share a considerable fraction of their adaptive evolution with CMC groups, specifically due to the putative adaptive genes described above and controlling intracellular Ca^2+^ mobilization, which exert crucial functions in both the *Glutamatergic synapse* and *Platelet activation* pathways (fig. 3d, supplementary table S5, Supplementary Material online). Nonetheless, all these loci turned out to be implicated also in immune and inflammatory responses, especially to cytomegalovirus and *T. cruzi*. Therefore, we can hypothesize that these (or similar) pathogens have plausibly represented the predominant selective pressures having acted on the ancestors of these Native Mexican populations, who have inhabited for millennia the coastal areas of southern Mexico, or the central-eastern coast in the case of the TOT. Indeed, these regions are characterized by a tropical savanna climate and represent the Mexican endemic areas of parasites such as *T. cruzi* and *L. Mexicana,* which are responsible for Chagas disease and localized cutaneous leishmaniasis (LCL) (RojoMedina et al. 2018; Loría-Cervera et al. 2019).

Interestingly, it has been demonstrated that platelets, besides playing a fundamental role in coagulation and vascular integrity, are also capable of controlling and modulating innate immune responses against pathogens, thus contributing to their clearance. This is mainly due to their abundance of membrane receptors, such as Tolllike receptors, which are able to detect pathogen- and damage-associated molecular patterns (Ribeiro, Migliari Branco, and Franklin 2019). In line with these findings and with our hypothesis of SMC adaptations to pathogens, a recent study identified three innate immune pathways, among which that of *Toll-like receptor signaling*, showing evidence of Native American ancestry enrichment polygenic adaptation in an Admixed

Mexican population of Mayan ancestry (Norris et al. 2020). In fact, platelet granule secretion and spread depend on increased intracellular Ca^2+^ concentration and the *ITPR2* gene that exhibited the most relevant selection signature and functional connection with the other putative adaptive loci in these Native Mexican populations (fig. 3d) is pivotal for mobilization of intracellular Ca^2+^ stores. Furthermore, calcium signaling is vital for host cell invasion and life cycle of both *T. cruzi* and *L. Mexicana* parasites. In particular, cell invasion by *T. cruzi* is Ca^2+^ dependent and its infective trypomastigote stages are capable of triggering intracellular free Ca^2+^ transients in its target cells through a mechanism involving the host IP3R and the TcIP3R homolog of *T. cruzi* (Docampo and Huang 2015; Hashimoto et al. 2016). Interestingly, infection by these parasites causes a peculiar clinical picture in southern Mexican populations. For example, in a 7-year longitudinal study conducted on subjects of southern Mexican ancestry chronically affected by Chagas disease, it was unexpectedly found that there was no progression of the main Chagas-related cardiomyopathies and no chagasic megaviscera compared to the chronic clinical picture shown by subjects from different regions of South America (Goldsmith et al. 1986). Similarly, variability of the clinical presentation of LCL has been observed, with populations from endemic areas often manifesting asymptomatic infection, suggesting a host-specific immune response capable of controlling parasite replication (Andrade-Narvaez et al. 2016). Taken together, these evidences and the findings obtained by gene network analyses performed on SMC populations suggest that they might have evolved adaptations in response to specific pathogens that are endemic to the regions long inhabited by their ancestors, possibly resulting in the enhancement of their innate immune-mediated mechanisms of clearance of infection.

### Conclusions

By imputing high-resolution genomic data for 271 individuals from 15 Mexican indigenous groups showing high proportions of NA ancestry, the present study succeeded in assembling the largest genomic dataset available so far for Native Mexican populations. This was crucial for investigating the as reliable as possible proxy of the full spectrum of Native Mexican variation, enabling us to explore patterns of fine-scale population structure among Mexican people and to reconstruct the unique adaptive histories of their ancestors. We thus provided new evidence for the considerably different selective pressures that have targeted the main genetic clusters identifiable within the overall Mexican population. Interestingly, some of the identified biological adaptations were found to have the potential to play a role in modulating susceptibility/resistance of these populations to certain pathological conditions. For instance, type 2 diabetes in Seri, impairment of gastrointestinal homeostasis after ingestion of mycotoxins in Rarámuri, dependence on addictive substances/foods in Central Mexican groups, and severity of Chagas-related comorbidities and infection by *L. Mexicana* in people from Southern Mexico. Accordingly, these findings represent valuable insights into the origins and the determinants of some peculiar biological traits of both Native and Mestizo Mexican people. Therefore, the obtained results might lay the foundation for future studies aimed at devising regionalized/personalized preventive strategies properly tailored to these human groups, which is considered the best course of action to cope with the health issues related to the ecological and cultural transitions that have been recently experienced by Mexican populations (Ojeda-Granados et al. 2017; 2020).

## Materials and Methods

### Analyzed Dataset

The data analyzed in the present study were retrieved from those constituting the NMDP, which were previously generated as described in Moreno-Estrada et al. (2014). This dataset included 350 samples from 15 Native Mexican populations and characterized for 903,800 single nucleotide polymorphisms (SNPs). In detail, two populations were from Northern Mexico (i.e. SER, TAR), six were from Central Mexico (i.e. HUI, NAH, NAJ, NAP, PUR, TOT), and seven were representative of people from Southern Mexico (i.e. MYA, LAC, TOJ, TZT, ZAP, MAZ, TRQ).

### Genotypes Imputation

To generate a high-resolution genomic dataset, the NMDP was used to perform genotype imputation. We first applied preliminary QC procedures using PLINK v2.0 (Chang et al. 2015) to retain only autosomal variants, as well as SNPs and samples showing genotyping success rate > 95%. This led to a filtered dataset made up of 347 samples characterized for 779,283 SNPs, whose genomic coordinates were re-annotated according to the human reference genome build 37 (hg19) using the *liftOver* tool (Hinrichs et al. 2006). We then phased this dataset by means of the Eagle v2.3 software (Loh et al. 2016) implemented in the Michigan Imputation Server (MIS) v1.0.4 (Das et al. 2016) and we used the MIS *Minimac3* algorithm to impute genotypes using the 1000 Genomes Project Phase 3 as reference panel (1000 Genomes Project Consortium et al. 2015). The imputed dataset was subsequently filtered to retain only high-quality genotypes by using the *bcftools* v1.8 package (Li et al. 2009) to remove variants showing a squared Pearson correlation coefficient (r^2^) ≤ 0.9 between the imputed genotype probability and the true genotype call, as suggested in Palmer and Pe’er (2016). This enabled us to obtain a highresolution dataset including 347 samples characterized for 6,375,183 SNVs.

### Data Curation

The imputed dataset was subjected to additional QC filters using PLINK v2.0 to retain SNVs and samples showing genotyping success rate > 95%, as well as variants with no deviations from the Hardy-Weinberg Equilibrium (*p*> 1.5×10^−9^ after Bonferroni correction for multiple testing). In addition, we filtered out SNVs with ambiguous A/T or G/C alleles and multiallelic variants.

The degree of recent shared ancestry (i.e. identity-by-descent, IBD) for each pair of subjects was also obtained by calculating the genome-wide proportion of shared alleles (i.e. identity by state, IBS) to remove closely related individuals. An IBD kinship coefficient (PI_HAT) ≥ 0.270 was thus considered to exclude 76 samples, which ultimately led to the generation of a “high-quality imputed dataset” that included information for 4,875,751 SNVs characterized on 271 individuals.

Genotype-based populations structure analyses were then used to check for consistency between the imputed and original NMDP data. PCA was computed using the *smartpca* method implemented in the EIGENSOFT package v6.0.1 (Patterson, Price, and Reich 2006), while the ADMIXTURE algorithm (Alexander, Novembre, and Lange 2009) was applied to test K = 2 to K = 12 population clusters by running 25 replicates with different random seeds for each K and by retaining only those presenting the highest log-likelihood values. In addition, crossvalidation errors were calculated for each K to identify the value that best fit with the data.

The “high-quality imputed dataset” was then phased using SHAPEIT2 v2.r790 (Delaneau et al. 2013) by applying default parameters and by using the HapMap phase 3 recombination maps and the 1000 Genomes Project dataset as a reference panel (1000 Genomes Project Consortium et al. 2015). Moreover, to perform haplotype-based selection scans, the “high-quality imputed dataset” was phased again with SHAPEIT2 v2.r790 (Delaneau et al. 2013) but considering information about ancestral and derived alleles as specified in the reconstructed reference human genome sequence (1000 Genomes Project Consortium et al. 2015).

### Haplotype-based Clustering Analyses

To identify groups of genetically homogenous Native Mexican populations and to remove potential outlier individuals not representative of their population of origin, we applied the haplotype sharing clustering methods implemented in the CHROMOPAINTER/fineSTRUCTURE package (Lawson et al. 2012) to the phased “high - quality imputed dataset”. In detail, the ChomoPainter v2 algorithm was used to produce a co-ancestry matrix through the reconstruction of the haplotype sharing pattern of each individual performed by considering all other subjects in the dataset as potential “donors” of DNA sequence chunks, but excluding themselves (Lawson et al. 2012). To this end, ten steps of the Expectation Maximisation algorithm were applied to a subset of chromosomes {4,10,15,22} to estimate recombination/switch rate and mutation/emission probability. The obtained values were averaged across the subset of chromosomes, by weighting them by the number of SNVs, and across individuals. The mean recombination/switch rate and mutation/emission probability values were then used to run again the ChomoPainter v2 algorithm on all autosomes by considering *k*= 50, which is a value preferable when examining closely related populations, to specify the number of expected haplotype chunks used to define a genomic region. We thus obtained a co-ancestry matrix by considering the matrix of counts of haplotype chunks shared among subjects, which were then summed across all autosomes. The obtained matrix was submitted to the Markov chain Monte Carlo (MCMC) fineSTRUCTURE v.fs2.1 algorithm (Lawson et al. 2012) by setting 1,000,000 burn-in iterations, followed by 1,000,000 more iterations and the sampling of inferred clustering patterns every 10,000 runs. A further fineSTRUCTURE v.fs2.1 run consisted in 100,000 additional hill-climbing iteration steps aimed at improving posterior probability and at merging the identified clusters in a step-wise fashion. The inferred haplotype sharing patterns were finally visualized as a circular dendrogram with the iTOL v5 tool (Letunic and Bork 2019), with clades of genetically homogenous populations being defined by considering only those branches of the dendrogram that were characterized by a posterior probability above 90%.

Pairwise F_st_ values between the identified population clusters were then computed according to the Weir and Cockerham method using the *4P* software (Benazzo et al. 2015). Calculation was replicated after pruning the dataset with PLINK v2.0 to maintain only SNVs in linkage equilibrium with each other and by using the functions implemented in the R *StAMPP* package. This enabled us to perform 1,000 bootstraps to obtain confidence intervals and corresponding *p*-values associated to the computed F_st_ values.

### Number of Segregating Sites by Length Statistic

Based on the assumption that the ancestors of Native Mexican people belonging to the same genetically homogeneous population cluster have plausibly shared a remarkable fraction of their evolutionary histories (Moreno-Estrada et al. 2014; Ávila-Arcos et al. 2020), having also experienced similar environmental- and/or cultural-related selective pressures, we searched for genomic signatures of natural selection independently in each population group.

For this purpose, we computed the nSL statistic, which was developed to detect a broad range of selective events ranging from strong positive selection on newly arisen alleles to more slight selection on multiple alleles, including standing variation (Ferrer-Admetlla et al. 2014). This improved the exploration of the full spectrum of adaptive events evolved by Native Mexican groups, including those mediated by genetic variants with a small individual effect size. In detail, nSL scores for each SNV were obtained through the application of algorithms implemented in the *selscan* v.1.1.0b program (Szpiech and Hernandez 2014) by setting a 20,000 bp threshold for gap scale, 200,000 bp as the maximum length a gap can reach, and 4,500 consecutive SNVs as the maximum extension parameter. The obtained unstandardized nSL scores were finally normalized across 100 frequency bins by subtracting to each value the average nSL score computed for the considered bin and by dividing the resulting score by the related standard deviation (Piras et al. 2012).

### Identifying Gene Networks Enriched for Selective Events

To test the occurrence of selective events under a realistic approximation of a polygenic adaptation model, we performed gene-network analyses by applying the *signet* algorithm implemented in the R package (Gouy, Daub, and Excoffier 2017). For this purpose, we used the genome-wide distribution of normalized nSL scores previously obtained for each population cluster to associate nSL values to each gene annotated on the human genome by considering SNVs within a range of 50 kb upstream and downstream of the chromosomal location of the genes. Then, we selected the highest nSL score among those computed for all variants ascribable to the same gene as the representative score of that given genetic locus. This *per gene* nSL value was used as input for the *signet* algorithm, which concomitantly considered also information available for all biological pathways annotated in the National Cancer Institute/Nature Pathway Interaction Database to assign the reconstructed gene networks to known functional pathways (Schaefer et al. 2009). In fact, the assumption beyond this approach was that multiple genes within a pathway and contributing to a certain biological function might interact to shape a given adaptive phenotype, thus being targeted simultaneously by natural selection. Therefore, by using a simulated annealing approach the *signet* algorithm searches for networks of genes that directly interact with each other and that at the same time present pervasive selection signatures (i.e. those whose distribution of scores within the pathway they belong to was shifted significantly towards extreme values). To this end, we set an iterative process of 20,000 iterations, and we generated null distributions of the highest scoring subnetwork (HSS) for each network of a specific size finally considering a significance threshold of 0.05 when comparing the obtained HSS with them. Accordingly, gene networks showing *p*-values < 0.05 were considered as significantly enriched for selective events occurred at multiple genes contributing to the same biological function and were plotted with Cytoscape v3.6.0 (Shannon et al. 2003). Moreover, we also performed a cross-check of the identified gene networks using the KEGG database (Kanehisa and Goto 2000) to identify additional pathways in which the putative adaptive genes might play an important role. Overall, this enabled us to shortlist the set of genes and functional pathways that have been more plausibly subjected to pervasive positive selection in the different population clusters.

## Data availability

The individual-level genotype data re-analyzed and imputed in the present study are available through a data access agreement by contacting the corresponding authors of the original source of the data as stated in Moreno-Estrada el al. 2014.

## Acknowledgments

We would like to thank all donors who contributed biological samples for the generation of the NMDP without whom this work would have not been possible. We are also grateful to Pier Massimo Zambonelli (CESIA, University of Bologna) for his IT assistance. C.O.G. was supported by two postdoctoral grants (grant n. 755288, 711247) from the National Council of Science and Technology (CONACyT) of Mexico in collaboration with the PhD Program in Molecular Biology in Medicine of the University of Guadalajara (CONACyT PNPC 000091) and the PhD Program in Integrative Biology, CINVESTAV, Iraputato (CONACyT PNPC 003649). S.D.S. was supported by the FONDAZIONE CASSA DI RISPARMIO IN BOLOGNA (grant n. 2019.0552 to M.S.).

